# Estimating Optimal Lysogenic Propensity for Viruses in Stressed Environments

**DOI:** 10.1101/321372

**Authors:** Devang Thakkar, Supreet Saini

## Abstract

*H*aving infected a bacterial cell, a temperate phage has to make a choice between (a) integrating itself into the bacterial genome, i.e. *lysogeny*, and (b) using the bacterial machinery to create multiple copies of itself and lysing the cell in the process, i.e. *lysis*. In order to maximize its long-term growth rate, phages need to ensure that they do not wipe off their bacterial hosts. Temperate phages have been observed to exhibit lysogenic propensities dependent on the *MoI* (Multiplicity of Infection), among other factors. We propose a model to estimate the propensity of lysogeny opted for by the phages in order to maximize coexistence. One possible approach to do so is to adopt a strategy that would help to attain and maintain an approximately equal proportion of phages with respect to their host. We find that the optimal fraction of phages opting for lysogeny follows a sigmoidal relationship with the *MoI* and is comparable to results obtained experimentally. We further assess the impact of phage and bacterial environmental stresses on the lysogenic propensity. Our results indicate that the optimal value of lysogenic propensity is greatly dependent on the intensity of these stresses.

## 1 Introduction

The primary aim of any organism is to avoid extinction, and to this end, it produces as many progenies as it is able to. In that case, ideally, bacteriophages must then opt only for lysis constantly without ever opting for lysogeny – an alternative that is less beneficial in the short run than the former. Such a policy is indeed the best strategy in the span of a few epochs, however the long term survival of the phages is dependent on their coexistence with their prey – the bacteria [1], [2], [3]. Temperate phages [4], [5] are phages that strike a balance between lysis and lysogeny in order to maximize long term survival. The lysogenic option makes it possible for phages to survive in the event of depletion of host bacterial cells. Previous studies on phage-host interaction have shown that the fraction of infected bacterial cells undergoing lysogeny is dependent on multiple factors including, but not limited to multiplicity of infection [6], [7] and environmental stresses [6], [8], [9]. Phages analyzed in these studies have been observed to opt for lysogeny more often when the multiplicity of infection is higher. The propensity of lysogeny is also reported to be higher when the environment is under stress, irrespective of whether the environment is lacking in required nutrients [6] or has excess concentrations of undesirable constituents [8].

Understanding the logic behind the lysis-lysogeny decision is important in order to comprehend how the phages are able to coexist with their hosts despite their high burst rates. A number of theoretical models [5], [10], [11], [12] have been proposed to explain the observed experimental trends. Although these models do provide a possible understanding of the system, they have their own limitations. Avlund et al [10] provide a game-theoretical model to explain the decision between lysogeny and lysis, the parameters for which are similar to those used in the current work. Avlund et al use the analogy of single player and multi-player games to explain the difference in results for *MoI* values of 1 and 2, and the use of discrete values of *MoI* is attributed to the fact that phages can assess only the number of phages in the cell they have infected. Sinha et al [11] model the effect of various factors – such as initial phage and bacterial populations, the burst size, the infection rate, and the bacterial growth rate – on the lysogenic propensity. The authors calculate the optimal lysogenic propensity by allowing different propensities to compete against each other under different system parameters. However, they do not consider the stresses on the environment which have been shown to affect the lysis-lysogeny decision. Also, the propensities are evolved over time for a few selected integral values of *MoI* (1, 2, 3) while leaving out the variation of lysogenic propensity with *MoI* at intermediate values. Maslov and Sneppen [5] model the interaction between phage and bacteria under environmental stresses using differential growth equations. One of the problems often faced by differential models for phage-bacteria interactions is that they require an infinite amount of bacteria because of the exponentially high growth rate of phages.

We attempt to resolve these issues and shortcomings in this study. Here, we use a single step simulation to estimate the optimal lysogenic propensity which sidesteps the infinite bacteria dilemma. Quorum sensing systems [13], [14] have been demonstrated in phages that allow them to communicate among themselves within and between generations. As a result phages can sense the number of phages not only within the cell, but also in the surrounding environment. This allows us to consider the estimation of the optimal propensity of lysogeny for non-integer values of *MoI* in our study. We propose below a single-epoch model that explains how a temperate phage maximizes its survival in the long run for varying environmental stresses and a continuous range of *MoI*. Our results indicate that this model is able to explain experimentally obtained trends of lysogenic propensity with respect to multiplicity of infection rather nicely.

## 2 Materials and Methods

Besides being characterized by the relative populations and growth rates, the phage-bacteria ecosystem implemented in our model is also dependent on the environmental stresses. We model the environmental stresses as the probability of ‘good’ and ‘bad’ environments for the two entities – the phage and the bacterium. Good environments favor the growth of the species while bad environments triggers a decay of the species, represented mathematically by an exponential decrease in the population. A good environment for the free phages is one wherein the phage population may successfully undergo lysis and increase the number of free phages. On the other hand, a bad free phage environment involves an exponential decay of the free phage population. Correspondingly, a good bacterial environment allows normal replicative growth of the bacterial population whereas a bad bacterial environment leads to an exponential decay of uninfected and lysogenized bacteria alike. The probability that an environment is good for the phage and bacterial population is denoted by *p*_1_ and *p*_2_ respectively. It then follows that the probabilities for a bad environment for phages and bacteria can be denoted by (1 *− p*_1_) and (1 *− p*_2_) respectively.

### 2.1 Assumptions

The relative population strength of the phages with respect to the bacteria is indicated by the *MoI*. The notion of *MoI* antedates the theory of phage infection commonly accepted today – that the adsorption of phages onto the bacteria follows a Poisson distribution [15]. Historically, *MoI* had been defined as the ratio of number of phages to the number of bacteria [16], [17]. However, over the years, various terminologies such as *MoIactual*, *MoIinput*, and *AP I (Average Phage Input)* have been coined to represent this quantity more correctly. For our discussion, we consider *MoI* to be the ratio of the effective number of phages (free phages + lysogenized phages) to the number of net bacteria (healthy + infected) in the system as explained in Equation 1.

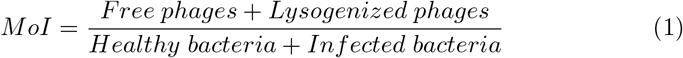

Secondly, we consider the infection of the bacteria by the phages to be quantified by a Poisson distribution with a mean equal to the *MoI*. This implies that for *N* bacteria that are exposed to phages at an *MoI* of *m*, *N *e^−m^* bacteria are not infected and would still be counted as healthy bacteria. Thirdly, we assume that for multiple infections of a bacterial cell, only one of the phages is effectively active [15]. In other words, super-infection of the host cell does not lead to any change in the state of the cell. A corollary of this is that the number of lysogenized phages can be considered to be equal to the number of infected bacteria. Following from Eq. 1, we thus get

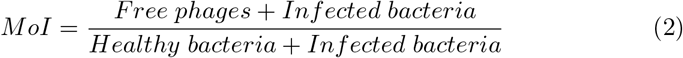

### 2.2 Parameters

We define below the different variables involved in our model and the Table 1 lists the notation and the values of the parameters of our model. As defined in Eq. 2, *MoI* is the ratio of the number of phages to the number of bacteria in the population. We choose the use the range covering the most commonly used *MoI* values and extend the range on both sides. We then divide the range into intervals of 0.01 to obtain a clearer understanding of the variation of lysogenic propensity as a function of *MoI*. The number of infected bacteria phages, and thereby the lysogenized phages (refer to Eq. 2) is denoted by *N_b,i_* whereas the number of healthy bacteria is denoted by *N_b,h_*. Since we start off with a completely uninfected bacterial population, the value of *N_b,i_* is set to 0 while *N_b_, h* is set to an arbitrarily large number. The free phage population size is denoted by *N_p,f_* whose initial magnitude is dependent on the *MoI* and the initial healthy bacterial population *N_b,h_* as

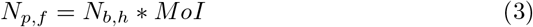

**Table 1.** List of parameters. This table lists out all the parameters in the model along with their symbol and values.

| Symbol | Parameter | Value(s) |
| --- | --- | --- |
| $MoI$ | Multiplicity of Infection | [0.01, 0.02, ..., 2] |
| $N_{b,h}$ | Number of uninfected bacteria | 10000 (Initial) |
| $N_{b,i}$ | Number of infected bacteria | 0 (Initial) |
| $N_{p,f}$ | Number of free phages | $N_{b,h} * MoI$ |
| $\gamma$ | Replicative growth rate | {1*, 2, 5} |
| $\alpha$ | Lytic burst rate | {10*, 20} |
| $p_1$ | Probability of good phage environment | [0.1, 0.2, ..., 1.0] |
| $p_2$ | Probability of good bacterial environment | [0.1, 0.2, ..., 1.0] |
| $\lambda_p$ | Phage decay rate in bad environments | {1, 2, 3} |
| $\lambda_b$ | Bacterial decay rate in bad environments | {0.1, 1} |
| $P_{lyso}$ | Lysogenic propensity | To be calculated |

(\* - Chosen)

The rate at which the species multiply is an essential parameter of a differential growth equation. We represent the phage burst rate by*γ* and the replicative growth rate of the bacterium by*α*. Values of *gamma* vary over a large range, usually within the range of 10–100 [15], but also going up to 300 in case of higher latent periods [18]. Since we are considering single step simulations, we define the value of*γ* as 10 in order to account for the latency period. Varying the relative magnitudes of these growth rates allows us to account for different system parameters such as the latency period and rate of infection.

### 2.3 The Model

Traditionally, stochastic simulations using the Gillespie Algorithm [19] have been carried out in order to understand the working of the lysis-lysogeny decision [20], [21]. The framework that researchers have worked with in the past requires them to consider an infinite supply of bacteria, while our method does not require us to do so.

Our model circumvents this problem by considering a single epoch simulation of the phage-bacteria interaction. We hypothesize that one way in which the phages may ensure their long-term survival is by striving for an approximately equal phage to bacterium ratio. In other words, the optimal strategy would be to choose the magnitude of the lysogenic propensity to be such that the resulting *MoI* is closest to unity. The rationale behind this is simple – if the *MoI* is less than 1, it means that the system still has “room” for the phage to grow more. On the other hand, if the *MoI* is already greater than 1, it means that there are probably few free bacteria that are yet to be infected. The interaction is modeled using the set of equations Eq. 4 to Eq. 15. The population dynamics of the two entities – the phages and the bacteria is modeled depending on the type of the external environment, thus giving us four different scenarios – good for both bacteria and phages, good for phages but bad for bacteria, bad for phages but good for bacteria, and bad for both phages and bacteria. In a good environment for the phages, the number of phages is multiplied by the burst size while in a bad environment for phages, the number of free phages decreases exponentially. Similarly, a good environment for bacteria sees the number of bacteria grow at the replicative growth rate, whereas a bad environment for bacteria leads to an exponential decay of the infected bacteria.

**Good phage environment, Good bacterial environment**

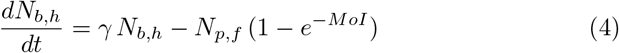

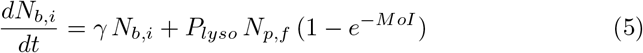

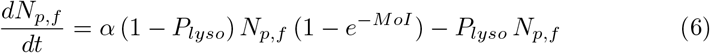

**Good phage environment, Bad bacterial environment**

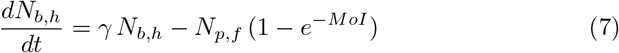

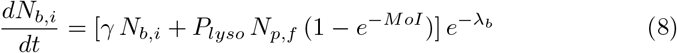

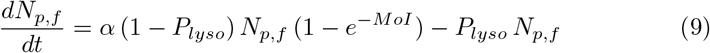

**Bad phage environment, Good bacterial environment**

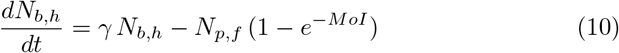

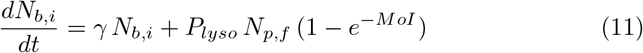

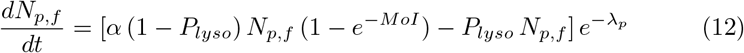

**Bad phage environment, Bad bacterial environment**

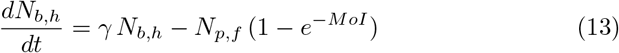

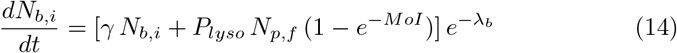

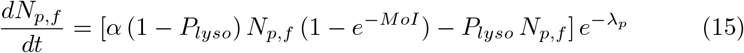

### 2.4 Simulation

Our aim here is to use the aforementioned equations and identify the optimal curve in the *P_lyso_* – *MoI* space for each value of the tuple (*p*_1_, *p*_2_) in the appropriate range.

1. For each value of the probabilities *p*_1_ and *p*_2_ indicated in Table 1, we establish a random phage and bacterial environment.
2. Next, we calculate the resulting *MoI* based on a single epoch resolution of the set of differential equations 4–15 for each value of initial *MoI* in the range {0.01, 0.02, …, 3}.
3. The value of *P_lyso_* that results an *MoI* closest to unity is chosen as the optimal value.

Steps 1–3 are repeated for 3000 iterations and the average value of the optimal *P_lyso_* is calculated. We used the Python programming language [22] for simulations and R [23] for creating the trellis plot.

## 3 Results

**The trend of the optimal lysogenic propensity**

The plot in Figure 1 shows the mean trend and maximum deviations of the optimal propensity of lysogeny for two extreme values of *p*_1_ as a function of *MoI* for variations in *p*_2_ (the probability of a good bacterial environment),*λ_b_* (the bacterial decay rate),*_p_* (the phage decay rate). The plot illustrates the robustness of the estimated *P_lyso_* for variation in the aforementioned parameters over the ranges mentioned in Table 1. As seen from Figure 1, the optimal strategy is to opt entirely for lysis as long as the relative phage concentration in the environment is lower than a threshold. Once the threshold is crossed, the value of *P* (*lyso*) increases rapidly and approaches one. The precise value of the *MoI* threshold is dependent on the quality of the environment, and the degradation rates for phages and bacteria.

**Fig. 1.**
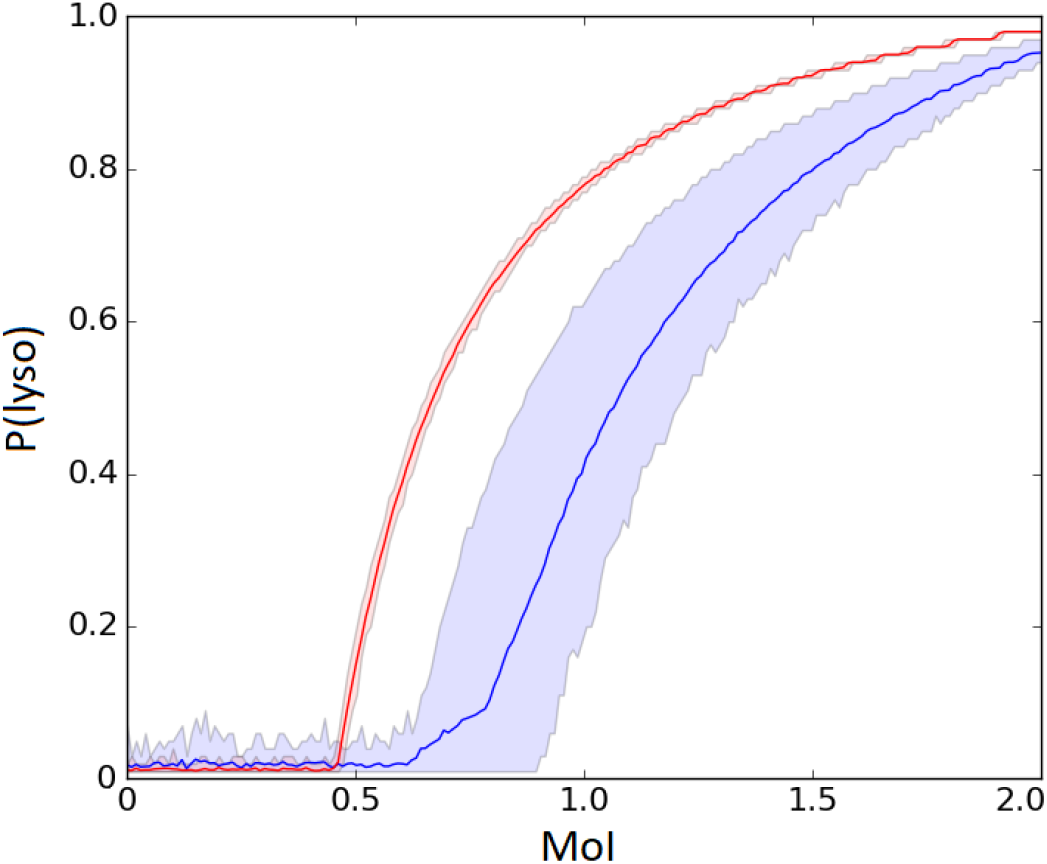
*P_lyso_* as a function of *MoI*. The blue curve refers to the mean values when *p*_1_ = 0.2, whereas the red curve refers to the mean values when *p*_1_ = 0.8; the shaded area represents the variation due to changes in the remaining parameters: *p*_2_ – (0.2, 0.4,0.6, 0.8),*λ_p_* – (1,2,3), and *λ_b_* – (0.1, 1). The curve in case of a good phage environment is more robust as compared to the bad phage environment where the other parameters play a significant role as well.

For bad phage environments, represented by the blue curve, the relation is affected more by changes in other parameters than good phage environments

8 D. Thakkar and S. Saini

- shown by the red curve with an extremely small error area. Classically, environments have been assumed to be good [11], [24], thus missing out on the variation caused by changes in the values of the parameters in bad environments. It is interesting to note that as the environment becomes worse for phages, the lysogenic propensity at a given *MoI* decreases. This follows from the fact that for an environment where the phages are rapidly dying, the phages need to produce a higher number of progeny in order to avoid being wiped off in the long run. Somewhat surprisingly, our simulations indicate that the change in the bacterial environment does not seem to affect the trend of lysogenic propensity as greatly as the change in the phage environment. Another factor which affects the *P_lyso_* versus *MoI* curve is the phage degradation factor*λ_P_*. As the value of_*p*_ increases, the curve shifts towards the left, with the shift being larger for bad phage environments and smaller for good phage environments. Figure 2 illustrates the relation between *P_lyso_* and *MoI* for a matrix of values of probabilities *p*_1_ and *p*_2_.

**Fig. 2.**
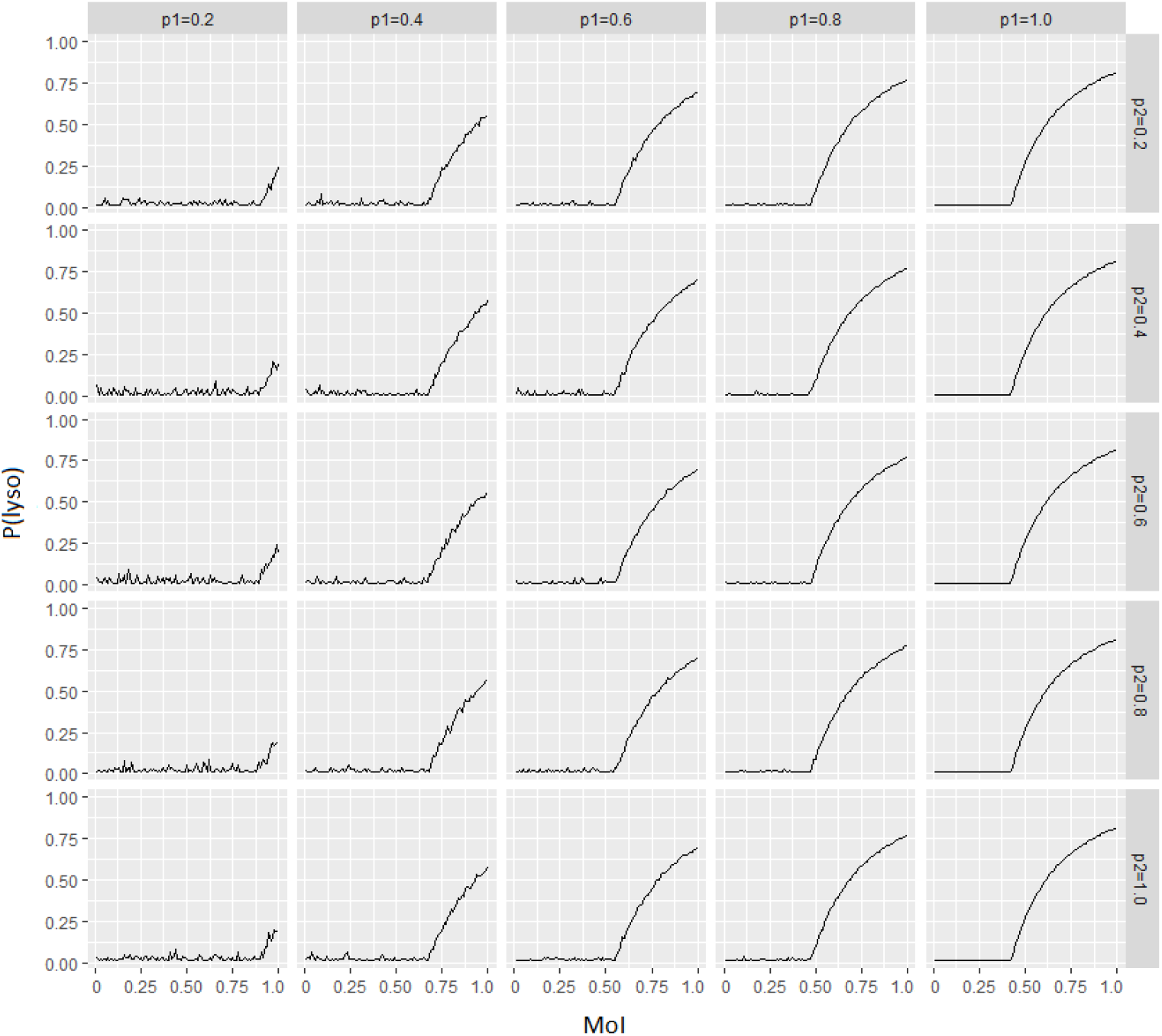
A Trellis plot displaying the variation in *P_lyso_* vs *MoI* trends as a function of *p*_1_ and *p*_2_. The foremost observation here is the variation of the curve as a function of *p*_1_. As the environment becomes better for phages, more and more phages opt for lysogeny. The change in the trend as the bacterial environments improve is subtler and is better perceived from the area under the graph.

Experimental research has shown that the lysis-lysogeny decision varies not only from species to species, but is also dependent on a variety of other factors including but not limited to multiplicity of infection, chemical environment, cell size, and location of inserted phage [6], [25], [26]. The problem that we try to address has been experimentally tested, albeit with the variation of different parameters [6], [26]. Our results match closely the results obtained in [26] and are qualitatively similar to the results presented in [6].

## 4 Discussion

In this work, we model the impact of environmental factors and the *MoI* on the optimal lysogenic propensity. There exists a gap in the literature on phage biology since how exactly the decision between lysis and lysogeny is made, and what range of factors impact this decision is currently not well understood. We look at the lysis versus lysogeny decision from the point of view of long-term coexistence using a single step simulation. In view of the fact that phages are indeed able to communicate the strength of their population [13] to future infecting phages, it is necessary to move away from the conventionally used integral *MoI* method. By selecting lysogenic propensities that lead to a resultant *MoI* of 1, we find that the results obtained match qualitatively with experimental data. A close matching to different experiments should be possible by using experimentally noted values for degradation, replication, and amplification rates. When the environment is more prone to bad episodes, the phages are more likely to opt for lysogeny. This can be seen as an example of bet-hedging, a concept that has been applied to the study of lysogeny in phages by [10], [27], [28]. Here, we evaluate the effects of various parameters considering bad phage and bacterial environments since restricting the study to good environments limits the observed variation due to change in parameters, as seen inFigure 1.

We have focused solely on how the phage lambda may survive in the long run in different environments and stresses, and how the optimal lysogenic propensity changes with the multiplicity of infection. In this work, we develop a model of estimating fraction of lysogeny that explains experimental data nicely. We have not addressed the question of how the gene regulatory network (GRN) of the phage can use the information about the environment to its benefit. It has been shown that the genetic switch in the phage can sense the concentration of bacterial substrates [29], so it might be able to use such a method to assess the environmental stress. Future work work would involve understanding the molecular underpinnings of how external information affects the gene network.

